# A kinase translocation reporter reveals real-time dynamics of ERK activity in Drosophila

**DOI:** 10.1101/2022.01.07.475336

**Authors:** Alice C. Yuen, Anadika R. Prasad, Vilaiwan M. Fernandes, Marc Amoyel

## Abstract

Extracellular Signal-Regulated Kinase (ERK) lies downstream of a core signalling cascade that controls all aspects of development and adult homeostasis. Recent developments have led to new tools to image and manipulate the pathway. However, visualising ERK activity *in vivo* with high temporal resolution remains a challenge in Drosophila. We adapted a kinase translocation reporter (KTR) for use in Drosophila, which shuttles out of the nucleus when phosphorylated by ERK. We show that ERK-KTR faithfully reports endogenous ERK signalling activity in developing and adult tissues, and that it responds to genetic perturbations upstream of ERK. Using ERK-KTR in time-lapse imaging, we made two novel observations: firstly, sustained hyperactivation of ERK by expression of dominant-active Epidermal Growth Factor Receptor raised the overall level but did not alter the kinetics of ERK activity; secondly, heterogeneity in ERK activity in retinal basal glia correlated with the direction of migration of individual cells. Our results show that KTR technology can be applied in Drosophila to monitor ERK activity in real-time and suggest that this modular tool can be further adapted to study other kinases.

**Summary Statement:** We describe a reporter to study the dynamics of ERK signalling in Drosophila, use it to measure signalling in individual cells over time, and monitor development.

## Introduction

Intercellular signalling patterns and instructs all aspects of development and adult homeostasis, from specifying cell fate to promoting morphogenesis, migration, and survival. As such, visualising where and when signalling occurs is critical to gaining a complete understanding of tissue formation and function. Visualising pathway activity often involves immunostaining for activated (often phosphorylated) components of the pathway, or using transcriptional reporters that are specifically regulated by the activity of the pathway. However, these approaches only provide snapshots of activity at particular timepoints, and do not capture the full extent of the dynamics of signalling. Indeed, the importance of temporal dynamics in signalling is increasingly being appreciated; for example, cells can integrate both the duration and amount of the Hedgehog morphogen they are exposed to and respond with differential gene expression (Dessaud et al., 2007). Similarly, although it had been known for many years that the Mitogen-Activated Protein Kinase (MAPK) pathway is required to specify the extraembryonic primitive endoderm in one of the first lineage restrictions in mammalian embryos (Chazaud et al., 2006), recent work showed that the levels of MAPK at mitotic exit, not at the point of fate acquisition, influence this fate decision (Pokrass et al., 2020).

Extracellular Signal-Regulated Kinase (ERK) is one of three MAPKs and plays critical and diverse roles throughout development. ERK is a serine-threonine kinase that is activated through phosphorylation and, in turn, affects diverse cellular processes, from gene expression to metabolism (Lavoie et al., 2020). Several receptor tyrosine kinases (RTKs) can initiate a phosphorylation cascade involving the GTPase Ras, and the kinases Raf and MEK, that ultimately leads to ERK phosphorylation and activation (Fig. 1A). One particularly notable feature of ERK signalling is the number of upstream receptors, and the fact that their activation of the same Ras-Raf-MEK-ERK cascade leads to different outcomes. For instance, in rat PC-12 cells, activation of the cascade via Epidermal Growth Factor (EGF) leads to cell proliferation, while stimulation with Nerve Growth Factor (NGF) results in differentiation (Greene and Tischler, 1976; Huff et al., 1981). Mounting evidence now shows that the quality and strength of upstream inputs are encoded as differential temporal kinetics of ERK activation, which are then decoded and read as instructive information to direct cellular responses (Albeck et al., 2013; Blum et al., 2019; Kholodenko et al., 2010; Marshall, 1995; Rauch et al., 2016; Ryu et al., 2015; Santos et al., 2007; Traverse et al., 1994). Indeed, exposure to repeated long pulses of low EGF can mimic sustained ERK activation typical of NGF stimulation and induce cell differentiation instead (Ryu et al., 2015). However, it is unknown whether information is encoded in the dynamics of signalling *in vivo*, where cells rarely encounter signals in such carefully timed pulses. This highlights the importance of studying ERK activity dynamics and the need for tools that allow visualisation of ERK kinetics of the same cells in a longitudinal manner.

**Figure 1.**
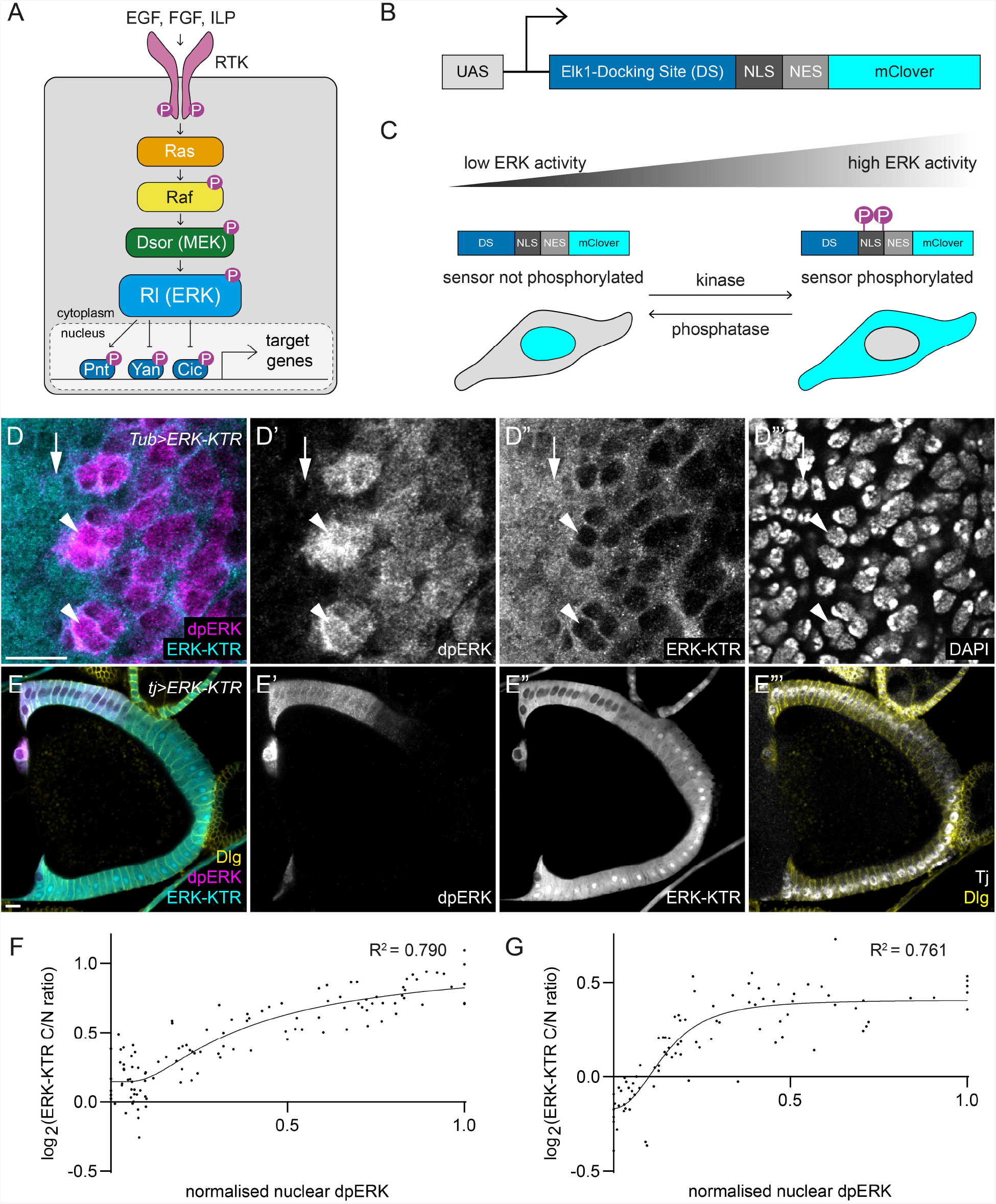
Drosophila ERK-KTR localisation changes with ERK activity in larval and adult tissues. (A) Schematic of ERK signalling in Drosophila. Ligands of the Epidermal Growth Factor (EGF), Fibroblast Growth Factor (FGF), and Insulin-Like Peptide (ILP) families activate their corresponding receptor tyrosine kinases (RTK) to trigger the activation of the canonical Ras-Raf-MEK-ERK phosphorylation cascade. In Drosophila, MEK is known as Dsor, and ERK is known as Rolled (Rl). Phosphorylation (represented by magenta circles labelled ‘P’) by ERK results in the activation of the transcriptional activator, Pointed (Pnt), and the inhibition of the transcriptional repressors, Yan and Capicua (Cic). (B) Drosophila ERK-KTR consists of the docking site (DS) for the mammalian ERK substrate and Pnt homologue, Elk1; a nuclear localisation signal (NLS); a nuclear export signal (NES); and the mClover green fluorescent protein, under the control of an Upstream Activator Sequence (UAS) promoter to enable tissue-specific expression of ERK-KTR in Drosophila. (C) ERK-KTR shuttles from nucleus to cytoplasm upon phosphorylation by ERK. Phosphorylation decreases the activity of the NLS and increases the activity of the NES, resulting in shuttling of ERK-KTR from the nucleus to the cytoplasm with increasing ERK activity. (D) ERK-KTR (cyan, single channel D”) expression in the eye imaginal disc. dpERK is shown in magenta (D’) and DAPI (D”‘) outlines cell nuclei. Arrowheads indicate cells with high dpERK expression, in which ERK-KTR is excluded from the nucleus. Arrow indicates a cell with low dpERK staining, showing nuclear enrichment of ERK-KTR. (E) ERK-KTR (cyan, single channel E”) expression driven in somatic follicle cells of an adult ovariole using the *tj*-*Gal4* driver. Follicle cell nuclei is labelled with Tj (white, E”‘) and membranes are labelled using Dlg (yellow, E”‘). dpERK (magenta, E’) in follicle cells is present in a gradient peaking at the anterior dorsal side of the egg chamber, which is reflected by a gradient of cytoplasmic to nuclear ERK-KTR (E”). (F,G) Quantification of ERK-KTR C/N ratio of the eye disc (F) and adult ovariole (G) showing positive correlation between the ERK-KTR C/N ratio and nuclear dpERK levels. dpERK was normalised to the minimum and maximum value for each sample, to account for variability in staining between samples. Both eye disc and ovariole data showed a good correlation when fitted to an asymmetric sigmoidal curve (R^2^ = 0.790 for eye discs, and R^2^ = 0.761 for ovarioles). Scale bar = 10 μm.

Many approaches have been taken to visualise ERK activity kinetics, including Förster resonance energy transfer (FRET) sensors, fluorescent fusion proteins for transcriptional effectors of the pathway, and phase separation-based kinase reporters (Lim et al., 2013; Moreno et al., 2019; Nakamura et al., 2021; Peláez et al., 2015; Zhang et al., 2018). One particularly promising approach is the development of a new class of kinase activity reporters, termed kinase translocation reporters (KTRs) (Regot et al., 2014; Spencer et al., 2013). KTRs consist of a docking site for a kinase-of-interest, fused to a fluorescent protein together with a nuclear localisation signal (NLS) and a nuclear export signal (NES). Both localisation signals are engineered to contain consensus phosphorylation sites for the kinase, such that phosphorylation decreases the activity of the NLS and increases the activity of the NES. Therefore, kinase activity is translated into shuttling of the chosen fluorescent protein from the nucleus to the cytoplasm, which can be tracked in live cells. One key advantage of the KTR method is its modularity, such that it can be easily implemented for many kinases, including ERK (Regot et al., 2014).

Given its genetic amenability, the fruit fly *Drosophila melanogaster* has proven a useful model to study ERK signalling. The Drosophila Ras-Raf-MEK-ERK cascade is conserved and activates the sole ERK, called Rolled (Rl) (Fig. 1A). Several RTKs lie upstream of ERK activity in flies, including the EGF Receptor (Egfr), two Fibroblast Growth Factor Receptors (Fgfr, encoded by the *heartless* and *breathless* loci), and the Insulin Receptor, to regulate cell behaviours including proliferation, growth, survival, differentiation, and morphogenesis (Hayashi and Ogura, 2020; Semaniuk et al., 2021; Shilo, 2014; Sopko and Perrimon, 2013). Importantly, the same complexity in the response to ERK signalling is seen as in other organisms; for instance, in somatic stem cells in the adult testes and ovaries, Egfr activity is required for both self-renewal and differentiation (Amoyel et al., 2016; Castanieto et al., 2014; Kiger et al., 2000; Schulz et al., 2002; Singh et al., 2016). During development, photoreceptors secrete both EGF and Fibroblast Growth Factor (FGF) ligands to signal to glial cells at the base of the eye imaginal disc. Egfr and Heartless in glia both signal through ERK to promote different outcomes: inducing differentiation of the photoreceptor target field and ensheathment of photoreceptor axons, respectively (Fernandes et al., 2017; Franzdóttir et al., 2009). Recent advances in the development of optogenetic tools to manipulate ERK signalling in flies have enabled a fine dissection of the temporal requirements for ERK activity during development, and refined our understanding of the signalling dynamics in cell fate decisions (Bunnag et al., 2020; Johnson and Toettcher, 2019; Johnson et al., 2017; Patel et al., 2019; Wang et al., 2020; Yadav et al., 2021). Thus, the ability to monitor ERK activity dynamics in Drosophila, in combination with the tools to manipulate it, would provide an unparalleled understanding of how temporal signalling dynamics regulate fate decisions.

Here we report a new KTR for ERK activity in Drosophila, named ERK-KTR, adapted from a previously published reporter developed in mammalian cells (Regot et al., 2014). We show that ERK-KTR reflects endogenous ERK signalling activity in both developing and adult tissues, and that it responds to genetic perturbations upstream of ERK. For ease of use in live imaging, we generated a reporter called ERK-nKTR, in which a red nuclear marker is used to segment and track cells, and to normalise nuclear ERK-KTR as a readout for kinase activity (de la Cova et al., 2017). We use this reporter to study ERK dynamics in the wing discs and find that overexpressing a dominant-active form of Egfr raised the overall level of ERK activation but did not result in altered patterns of activity. Finally, we demonstrate the use of this reporter in migrating cells by examining ERK activity in glia in the optic stalk. We found that ERK activity correlated with the direction of migration, indicating that ERK levels distinguish between naïve glia and those that have been exposed to photoreceptor-derived signals. In sum, we show that KTR technology can be applied in Drosophila to monitor ERK levels, suggesting that its modular design could be altered to study the dynamics of multiple other kinases.

## Results

### ERK-KTR localisation reflects endogenous patterns of ERK activity

To monitor ERK activity *in vivo*, we adapted a previously published ERK-KTR (Regot et al., 2014) for use in Drosophila. This reporter consists of the ERK docking site from the mammalian ERK substrate, Elk1, fused to mClover with phosphorylation-sensitive NLS and NES sequences (Fig. 1B,C). Upon phosphorylation by ERK, the activity of the NLS is reduced while that of the NES is enhanced, resulting in the shuttling of the KTR from the nucleus to the cytoplasm. Therefore, a high cytoplasmic-to-nuclear (C/N) fluorescence ratio should reflect high ERK activity (Fig. 1C). We generated flies carrying a transgenic construct encoding ERK-KTR under the control of an Upstream Activator Sequence (UAS) promoter, such that the expression of ERK-KTR can be driven by Gal4 in a tissue-specific manner.

To validate that this construct reports endogenous ERK activity, we analysed larval and adult tissues in which the patterns of ERK activity are well-characterised. We expressed ERK-KTR under the control of *Tubulin* (*Tub*)-*Gal4* and performed immunofluorescence against doubly phosphorylated ERK (dpERK) to detect endogenous, active ERK. In third instar larval eye-antennal imaginal discs, ERK activity is required for differentiation of photoreceptors (Roignant and Treisman, 2009). A band of apical constriction, called the morphogenetic furrow, progressively sweeps across the eye disc from posterior to anterior. Posterior to the furrow, founding photoreceptors express the EGF, Spitz, leading to ERK activation in neighbouring cells and their recruitment into ommatidial clusters. We examined ERK-KTR subcellular localisation in third instar larval eye discs (Fig. 1D). In clusters of cells expressing high levels of dpERK, ERK-KTR was mostly cytoplasmic and excluded from the nucleus (identified by DAPI staining; Fig. 1D”, arrowheads). In contrast, in neighbouring cells with low dpERK levels, ERK-KTR was present in higher levels in the nucleus (Fig. 1D”, arrow). To quantify the C/N ratio of ERK-KTR, we measured mClover levels in the nucleus and in a 3 pixel-wide perinuclear ring. These measurements showed that the C/N ratio of ERK-KTR was positively correlated with dpERK levels (Fig. 1F, R^2^ = 0.790, N = 116 cells across 5 discs). Thus, ERK-KTR localisation is a good readout for dpERK levels in the eye disc.

Having shown that ERK-KTR can be used to monitor ERK activity in the larva, we asked whether the same was true in adult tissues. In the ovariole, germline cysts are surrounded by an epithelium composed of somatic follicle cells (Wu et al., 2008). At stage 9/10a, expression of the Gurken ligand by the oocyte produces a gradient of ERK activity at the dorsal anterior side of the follicular epithelium (Gonzâlez-Reyes et al., 1995; Peri et al., 1999; Wasserman and Freeman, 1998; Fig. 1E’). We used *traffic jam* (*tj*)-*Gal4* to drive ERK-KTR expression specifically in follicle cells. We observed that mClover was excluded from the nucleus of dorsal anterior follicle cells, gradually becoming more enriched in the nucleus in more posterior and ventral cells, forming a gradient of nuclear exclusion to nuclear enrichment (Fig. 1E”). We quantified the ERK-KTR C/N ratio in follicle cells using Tj immunofluorescence to identify the nucleus and Discs Large (Dlg) to identify cell outlines and measure cytoplasmic intensity (Fig. 1E”). As in the eye disc, we found a strong positive correlation between the ERK-KTR C/N ratio and dpERK levels in follicle cells (Fig. 1G, R^2^ = 0.761, N = 90 cells in 5 ovarioles).

We observed that, depending on the driver used, expression levels of the ERK-KTR could be variable from cell to cell. We sought to determine whether the level of ERK-KTR expression impacted the C/N ratio of ERK-KTR. However, there was no correlation between total ERK-KTR expression and the ERK-KTR C/N ratio (Fig. S1A, R^2^ = 0.034 for eye disc; Fig. S1B, R^2^ = 0.003 for ovariole). Thus, the ERK-KTR C/N ratio accurately reflects endogenous ERK activity levels in both larval and adult tissues, independently of the overall expression level.

### ERK-KTR localisation responds to genetic perturbations in the ERK signalling pathway

Having shown that ERK-KTR localisation correlates with endogenous ERK activity, we next asked whether it would respond to genetic manipulations of the ERK pathway. To avoid confounding interpretations of our results due to endogenous patterns of ERK activity, we turned to the third instar larval wing disc, where there are low, uniform levels of ERK activity. We generated flip-out clones in the wing disc pouch that expressed ERK-KTR and quantified the ERK-KTR C/N ratio in tissue cross-sections where the nucleus and the cytoplasm could be outlined using DAPI and Dlg staining, respectively (Fig. 2A-H). Control flip-out clones (Fig. 2A,B) expressing ERK-KTR alone in the wing disc pouch showed mixed cytoplasmic and nuclear localisation of mClover. In contrast, ERK-KTR was excluded from the nucleus in clones expressing a constitutively active form of Ras, Ras^V12^ (Fig. 2C,D). Conversely, clonal expression of a dominant-negative form of Egfr (DN-Egfr) resulted in increased nuclear localisation of mClover (Fig. 2E,F). Expression of Ras^V12^ resulted in a significantly higher ERK-KTR C/N ratio than in control clones (Fig. 2I, P<0.0032, one-way ANOVA followed by Holm-Šidák’s multiple comparisons test, N = 42 cells across 6 discs for control, N = 38 cells across 6 discs for Ras^V12^), while expression of DN-Egfr led to a lower ratio (Fig. 2I, P<0.0124, N = 46 cells across 5 discs). We also tested whether other kinases could alter ERK-KTR localisation by inducing clones overexpressing the Drosophila Janus kinase, Hopscotch (Hop). As expected, we did not observe any significant changes in ERK-KTR localisation compared to control (Fig. 2G-I, P<0.9984, N = 43 cells across 4 discs). Thus, ERK-KTR localisation responds specifically to manipulations that affect ERK signalling.

**Figure 2.**
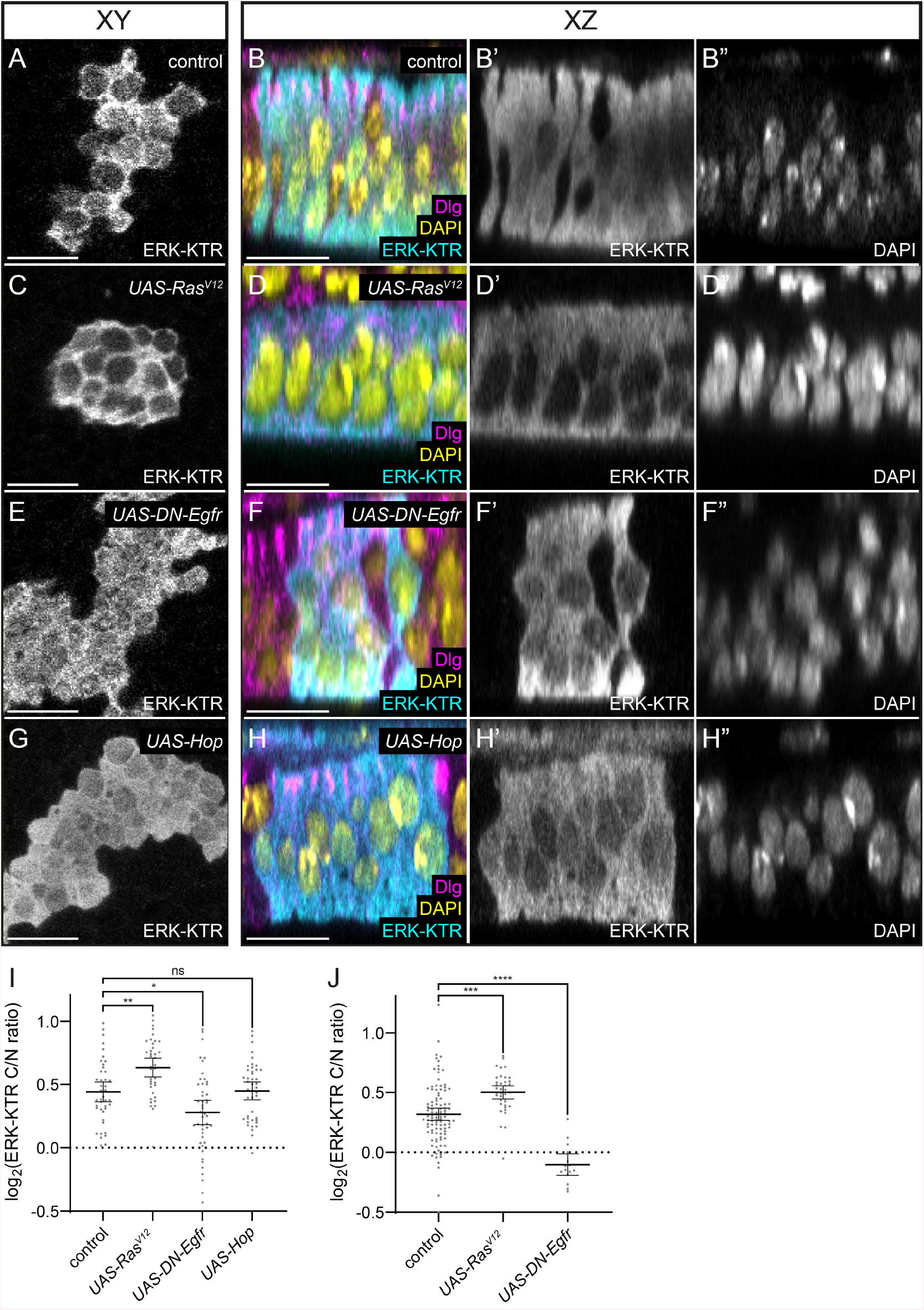
Genetic perturbation of the ERK pathway results in changes to ERK-KTR localisation. (A-H) Wing disc clones expressing Drosophila ERK-KTR, shown in the plane of the epithelium (A,C,E,G) or in cross-section (B,D,F,H). ERK-KTR is shown in cyan (B,D,F,H) and single channels are seen in A,C,E,G and B’,D’,F’,H’. Dlg is shown in magenta and DAPI in yellow (single channel B”,D”,F”,H”). In control clones (A,B), ERK-KTR localisation varied among cells within the clone with some cells displaying nuclear localisation, and others predominantly cytoplasmic localisation. (C,D) Clones expressing a constitutive-active form of Ras, Ras^V12^, in which ERK signalling was hyperactivated, showed nuclear exclusion of ERK-KTR. (E,F) Expression of a dominant-negative form of Egfr (DN-Egfr) resulted in increased nuclear localisation of ERK-KTR. (G,H) Overexpression of Hopscotch (Hop), a kinase that is not in the ERK signalling pathway, did not alter ERK-KTR localisation compared to control. (I) Quantification of the ERK-KTR C/N ratio in clones of the indicated genotype. Localisation was significantly affected by Ras^V12^ expression (P<0.0032, one-way ANOVA followed by Holm-Šidák’s multiple comparisons test, N = 42 cells across 6 discs for control, N = 38 cells across 6 discs for Ras^V12^) and DN-Egfr expression (P<0.0124, N = 46 cells across 5 discs), but not Hop overexpression (P<0.9984, N = 43 cells across 4 discs). (J) Quantification of the ERK-KTR C/N ratio in the somatic cyst lineage of adult testes. ERK-KTR was expressed in cyst stem cells and early cyst cells using the *C587*-*Gal4* driver either alone (control) or together with Ras^V12^ or DN-Egfr. Compared to control, Ras^V12^ expression significantly increase the ERK-KTR C/N ratio (P=0.0001, one-way ANOVA followed by Holm-Šidák’s multiple comparisons test, N = 102 cells across 20 testes for control, N = 38 cells across 5 testes for Ras^V12^), while DN-Egfr significantly decreased it (P<0.0001, N = 15 cells across 4 testes). Bars show mean and 95% confidence interval. ns: not significant. Scale bar = 10 μm.

Next, we used *C587*-*Gal4* to express ERK-KTR in the somatic cyst lineage of adult testes to assess how ERK-KTR localisation was affected by manipulating ERK activity in adult cells. We quantified ERK-KTR C/N ratio in the somatic cyst stem cells using Dlg immunofluorescence to outline the membrane and the nuclear dye, To-Pro-3, to identify the nucleus. Similar to our observations in the wing disc, Ras^V12^ expression resulted in a significant increase in the ERK-KTR C/N ratio compared to control (Fig. 2J, P=0.0001, one-way ANOVA followed by Holm-Šidák’s multiple comparisons test, N = 102 cells across 20 testes for control, N = 38 cells across 5 testes for Ras^V12^), while DN-Egfr expression led to a significant decrease in the ratio (Fig. 2J, P<0.0001, N = 15 cells across 4 testes).

These data indicate that manipulating ERK activity results in change in ERK-KTR localisation, suggesting that, as in vertebrates, ERK-KTR does indeed directly interact with ERK through its docking site from the mammalian Elk1 (Yang et al., 1998). Since the reporter is expressed at non-physiological high levels, we asked whether it could quench endogenous ERK, leading to defects in ERK signalling. To test whether this was the case, we examined the pattern of veins in adult wings, which is acutely sensitive to ERK activity during development (Brunner et al., 1994). Patterning of wing veins was not affected by expressing ERK-KTR ubiquitously under the control of *Tub*-*Gal4* (Fig. S2A,B, N = 36 wings for *Tub*-*Gal4* control, N = 45 wings for *Tub>ERK-KTR*). Moreover, we observed no significant difference in the size of the wings between control and ERK-KTR-expressing animals (Fig. S2C, P<0.0743, Student’s t-test), indicating that growth was not affected. Altogether, our results show that ERK-KTR specifically responds to changes in ERK activity in both larval and adult tissues, and does not interfere with endogenous signalling.

### A modified ERK-KTR enables ERK activity measurements from nuclear fluorescence alone

Although the intended readout of the ERK-KTR is the C/N ratio, this measurement may be difficult to obtain in time-lapse live imaging experiments where both nucleus and cell outlines may not be labelled, or for cells with irregular morphologies, such as neurons or glia. Therefore, we sought to determine whether the intensity of nuclear mClover alone could be used to measure ERK activity, as a proxy for the C/N ratio of ERK-KTR. In the experiments described above (Fig. 2A-I), Ras^V12^ overexpression resulted in an enrichment in ERK-KTR localisation to the cytoplasm, which was clearly visible and significantly different from control when ERK-KTR C/N ratio was quantified (Fig. 2I). However, measuring only the nuclear ERK-KTR intensity did not reveal a significant change from control (Fig. S3A, P=0.997, one-way ANOVA followed by Holm-Šidák’s multiple comparisons test), while Hop overexpression resulted in a significant reduction (Fig. S3A, P<0.0001). Importantly, we observed significant differences in total (nuclear + cytoplasmic) ERK-KTR expression levels among the clones of different genotypes (Fig. S3B), suggesting that the variability in expression level prevents direct comparison of nuclear levels. Thus, measuring nuclear ERK-KTR levels alone is not sufficient for accurate assessment of ERK activity, and it is important to normalise for ERK-KTR expression differences by using the C/N ratio.

To overcome this issue and to develop a reporter that could be useful in live imaging, we adapted the method developed by de la Cova *et al*. (2017), termed ERK-nuclear KTR (ERK-nKTR), allowing for ERK-KTR measurements using just nuclear intensities. ERK-nKTR encodes ERK-KTR and a Histone 2Av (His2Av)-mCherry fusion, separated by a self-cleaving T2A peptide (Fig. 3A). Since they are translated as one peptide, the red and green fluorophores are produced in equimolar amounts, enabling normalisation of the total ERK-KTR expression levels; here, increased ERK activity should be reflected in an increase in red-to-green (R/G) ratio (Fig. 3B). Additionally, we generated non-phosphorylatable and phospho-mimetic variants of ERK-nKTR to determine the dynamic range of the reporter.

**Figure 3.**
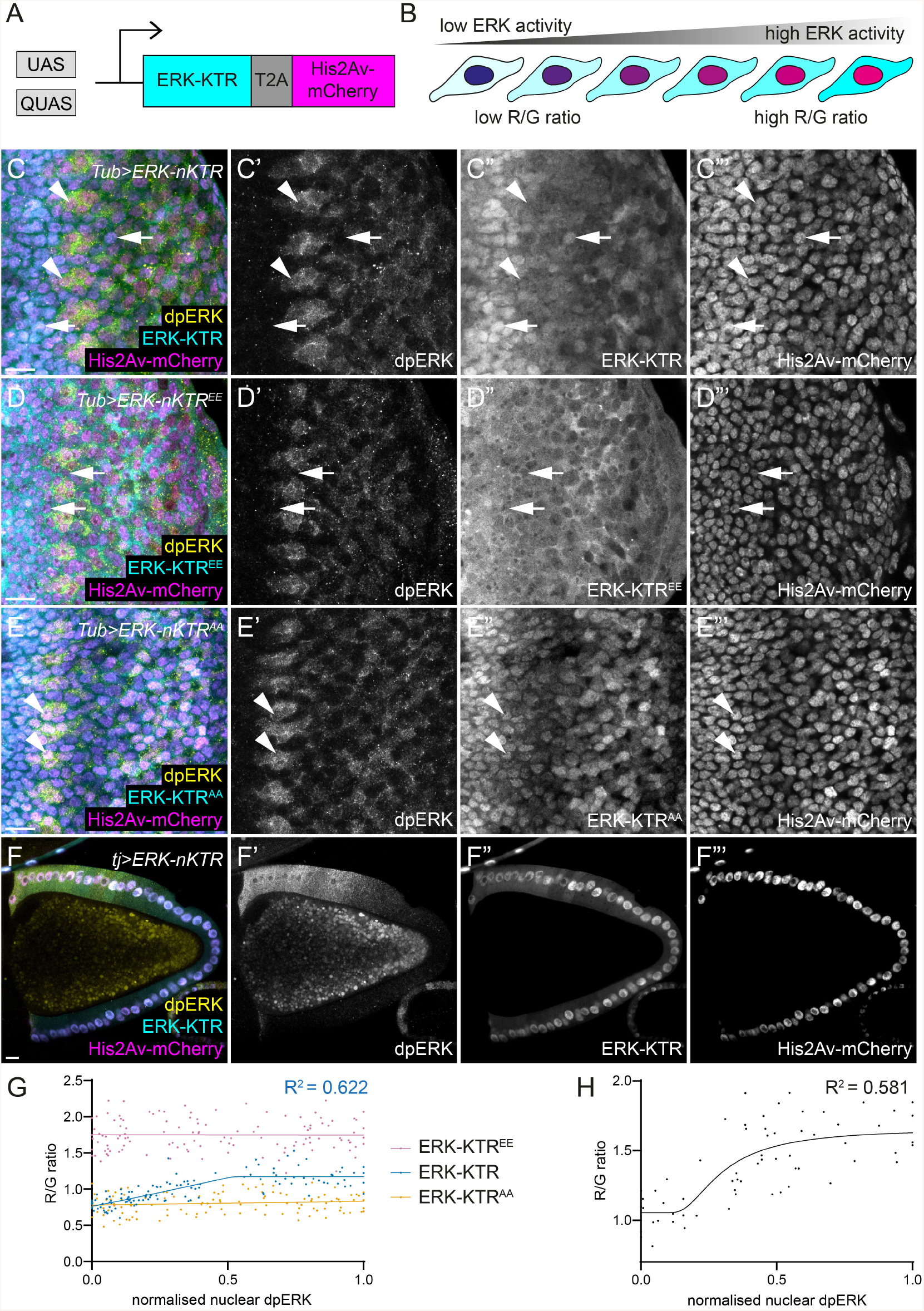
ERK-nKTR allows measurement of ERK activity using nuclear fluorescence intensity. (A) Schematic of the Drosophila ERK-nKTR. The Gal4/UAS and Q systems can be used to express ERK-KTR together with a Histone 2Av (His2Av)-mCherry fusion protein, encoded in a single open reading frame and separated by a self-cleaving T2A peptide. Red and green fluorophores are produced in equimolar amounts to enable normalisation of the total ERK-KTR expression levels. (B) Increased ERK activity results in nuclear exclusion of ERK-KTR and therefore an increase in the red-to-green (R/G) ratio. Using cyan and magenta, this results in a blue nucleus with low ERK activity and a magenta nucleus with high ERK. (C-F) ERK-KTR is shown in cyan and single channels in C”-F”, His2Av-mCherry is shown in magenta and C”‘-F”‘, and dpERK is shown in yellow and in C’-F’. (C) Subcellular localisation of ERK-nKTR (cyan, C”) expressed in the eye imaginal disc with *Tub*-*Gal4* follows the pattern of endogenous ERK activity, detected by labelling with an anti-dpERK antibody. Arrowheads indicate cells with high dpERK staining, in which ERK-nKTR is predominantly cytoplasmic, resulting in nuclei with a more magenta appearance in overlay due to the presence of mCherry-tagged His2Av. Arrows point to cells with low dpERK, where ERK-nKTR is enriched in the nucleus and the nuclei are blue in the overlay with His2Av-mCherry. (D) A phospho-mimetic variant of ERK-KTR, ERK-KTR^EE^, is constitutively located in the cytoplasm, regardless of endogenous ERK activity levels. Arrows indicate cells with low dpERK where ERK-KTR^EE^ is excluded from the nucleus, resulting in a magenta appearance of the nuclei in the overlay with His2Av-mCherry. (E) A non-phosphorylatable variant of ERK-KTR, ERK-KTR^AA^, is constitutively located in the nucleus, regardless of endogenous ERK activity levels. Arrowheads indicate cells with high dpERK, where ERK-KTR^AA^ is enriched in the nucleus, resulting in the blue appearance of the nuclei in the overlay with His2Av-mCherry. (F) ERK-nKTR expressed in somatic follicle cells of an adult ovariole with *tj*-*Gal4*. At this stage, dpERK is distributed in a gradient from the anterior dorsal side. ERK-KTR is localised in a matching gradient from cytoplasmic enrichment to nuclear enrichment, resulting in a magenta-to-blue gradient of nuclei in the overlay with His2Av-mCherry. (G) Quantification of R/G ratios of ERK-nKTR, ERK-nKTR^EE^, and ERK-nKTR^AA^ in the eye imaginal disc. ERK-nKTR R/G ratio as a function of normalised nuclear dpERK levels was fitted to an asymmetric sigmoidal curve and showed a positive correlation (blue line, R^2^ = 0.622). ERK-nKTR^EE^ and ERK-nKTR^AA^ R/G ratios as a function of normalised dpERK levels were fitted to simple linear regressions (pink and yellow, respectively). (H) Quantification of ERK-nKTR R/G ratio in somatic follicle cells. ERK-nKTR R/G ratio as a function of normalised nuclear dpERK levels was fitted to an asymmetric sigmoidal curve (R^2^ = 0.581). Scale bar = 10 μm.

To test these constructs, we expressed them ubiquitously using *Tub*-*Gal4* and dissected eye discs and adult ovarioles, as above. In the eye disc, ERK-nKTR appeared to be mostly localised to the cytoplasm in cells with high dpERK (Fig. 3C, arrowheads), and to the nucleus where dpERK intensity was low (Fig. 3C, arrows). In contrast, the phospho-mimetic ERK-KTR^EE^ was constitutively located in the cytoplasm (Fig. 3D, arrows indicate ERK-KTR^EE^ cytoplasmic localisation in cells with low dpERK), while the non-phosphorylatable ERK-KTR^AA^ was enriched in the nucleus (Fig. 3E, arrowheads indicate ERK-KTR^AA^ nuclear enrichment in cells with high dpERK), irrespective of dpERK levels. We quantified the R/G ratio of ERK-nKTR in the eye disc and found that the R/G ratio increased with dpERK levels (Fig. 3G, R^2^ = 0.622, N = 120 cells across 4 discs), in a manner comparable to the ERK-KTR C/N ratio of the original version of the reporter (Fig. 1F,G). Importantly, the R/G ratios of ERK-nKTR^AA^ and of ERK-nKTR^EE^ were constant and independent of dpERK levels (Fig. 3G, N = 119 cells across 4 discs for ERK-nKTR^AA^, N = 120 cells across 4 discs for ERK-nKTR^EE^); they represented the minimum and maximum of the dynamic range of the reporter, respectively, in this set of experiments. Finally, we used *tj*-*Gal4* to express ERK-nKTR in the ovary (Fig. 3F) and observed a similar positive correlation of the ERK-nKTR R/G ratio with dpERK (Fig. 3H, R^2^ = 0.581, N = 69 cells across 4 ovarioles).

In sum, ERK-nKTR is an effective phosphorylation-dependent reporter for the ERK pathway, enabling simpler measurements of ERK activity using only nuclear fluorescence intensities.

### ERK dynamics upon Egfr gain-of-function in time-lapse live imaging experiments

Having shown that ERK-nKTR can be used to monitor ERK levels in fixed tissue, we asked whether we could detect temporal patterns of ERK activity using ERK-nKTR in time-lapse movies. Previous studies in cell culture have shown that different kinetics of ERK activity can lead to different responses, such as proliferation or differentiation (Marshall, 1995). However, *in vivo* manipulations tend to involve permanent or prolonged activation of the pathway – expression of a dominant-active construct, for instance. Thus, we asked what effect hyperactivating the upstream receptor, Egfr, had on the dynamics of ERK signalling *in vivo*.

We induced control flip-out clones expressing ERK-nKTR in the early third instar larval wing disc (Fig. 4A and Movie S1) or clones expressing both ERK-nKTR and a dominant-active form of Egfr, λTop (Fig. 4B and Movie S2). At this developmental stage, there is no discernible spatial pattern of ERK activity, allowing us to compare clones across the wing pouch region. Explanted discs were embedded in agarose gel and imaged at two-minute intervals for two hours. In λTop-expressing clones, ERK-KTR was visibly more cytoplasmic than in control clones (Fig. 4A,B, arrowheads), consistent with higher ERK activity. Indeed, when we stained fixed discs with an antibody against dpERK, λTop-expressing clones showed high dpERK levels (Fig. S4A,B).

**Figure 4.**
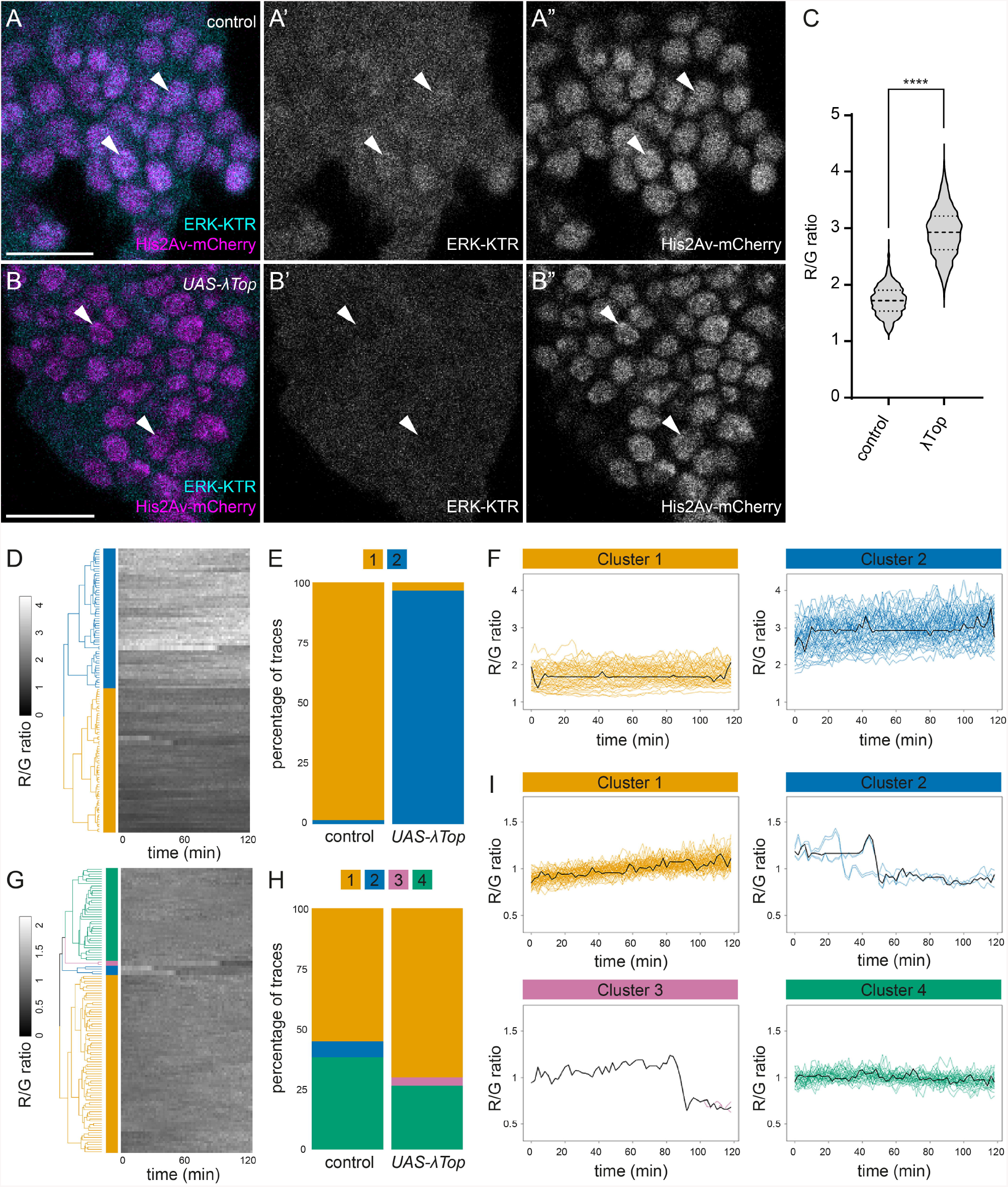
Egfr gain-of-function raises overall level of ERK activity without altering ERK kinetics. (A,B) Stills from movies of clones in third instar wing imaginal discs expressing ERK-nKTR with ERK-KTR in cyan (A’,B’) and His2Av-mCherry in magenta (A”,B”). Control clones (A) showed cells with nuclear enrichment of ERK-KTR (arrowheads). Clones expressing a dominant-active form of Egfr (λTop) (B) showed lower nuclear localisation of ERK-KTR (arrowheads). (C) Violin plot of all the ERK-nKTR R/G ratio measurements for control and λTop-expressing clones (N = 60 timepoints for 60 cells from 3 movies for each genotype). The mean ERK-nKTR R/G ratio in control clones was significantly lower than in λTop-expressing clones (1.72 ± 0.26 for control, 2.92 ± 0.43 for λTop, P<0.0001, Student’s t-test). Thick dotted lines indicate the median and thin dotted lines indicate upper and lower quartiles. (D) Heatmap showing the ERK-nKTR R/G ratio traces over time with each row corresponding to a single-cell. Traces were hierarchically clustered with dynamic time warping. The resulting dendrogram is shown on the left, where two main clusters were identified, colour-coded in blue (Cluster 1) and yellow (Cluster 2). (E) Bar chart showing the percentage of traces falling within each cluster for control and λTop-expressing cells. Clusters are represented using the colour-coding shown in panel D. (F) Plots showing the traces for ERK-nKTR R/G ratios from the two clusters, using the colour-coding shown in panel D, and the cluster average trajectories (black line). (G) Heatmap showing ERK-nKTR R/G traces and dendrogram resulting from performing hierarchical clustering with dynamic time warping on traces that have been normalised to its own mean to remove intensity differences. Four clusters were identified, colour-coded in yellow (Cluster 1), blue (Cluster 2), pink (Cluster 3), and green (Cluster 4). (H) Bar chart showing the percentage of traces falling within each cluster for control and λTop-expressing cells, using the same colour code as in panel G. (I) Plots showing traces for ERK-nKTR R/G ratios from each cluster, using the colour code shown in panel G, and the cluster average trajectories (black line). Cluster 2 and Cluster 3 correspond to cells that underwent division during the time-lapse movies. Scale bar = 10 μm.

To determine the dynamics of ERK activity in these clones, we manually segmented nuclei using the His2Av-mCherry channel and measured the ERK-nKTR R/G ratios in 20 cells per clone over time (N = 3 movies per genotype). Consistent with our qualitative observations, the mean ERK-nKTR R/G ratio in λTop-expressing clones was higher than in control clones (Fig. 4C, 1.72 ± 0.26 for control vs. 2.92 ± 0.43 for λTop, P<0.0001, Student’s t-test). Next, we sought to determine whether any patterns of activity could be detected across the control and hyperactivation conditions. We performed hierarchical clustering with dynamic time warping on ERK-nKTR R/G ratio traces pooled from both conditions to extract shape features that may not be aligned in time between traces (Blum et al., 2019; Giorgino, 2009). This analysis grouped all the traces into two clusters (Figs 4D-F and S5A), which largely segregated the control and λTop-expressing cells (Fig. 4E). However, there was no discernible pattern to the ERK-nKTR R/G ratio traces, which remained mostly steady in both clusters (Fig. 4F), suggesting that the difference between these clusters was largely due to the absolute values of ERK-nKTR R/G ratios.

In order to test this possibility, we rescaled each trajectory around its mean prior to running the clustering algorithm, such that information about the absolute value was removed and only changes in the ratio over time were analysed. In this case, clustering the traces resulted in four clusters (Figs 4G-I and S5B). The majority of trajectories fell into either Cluster 1, which was composed of traces showing a gentle upward trend of ERK-nKTR R/G ratios (Fig. 4I, yellow traces), or Cluster 4, which was composed of traces with steady ERK-nKTR R/G ratios (Fig. 4I, green traces). These clusters contained both control and λTop-expressing cells in similar proportions (Fig. 4H). Thus, the ERK dynamics of control and of λTop-expressing cells can no longer be distinguished if information about absolute levels of activity is removed. Few trajectories formed Cluster 3 and Cluster 4 (Fig. 4H); these traces corresponded to cells that had divided during the time-lapse movies. This is consistent with previous reports of a characteristic rise in ERK-KTR C/N ratio just prior to division and a drastic drop upon cytokinesis, followed by a recovery phase (Fig. 4I, blue traces and pink traces) (Pokrass et al., 2020). In conclusion, we have successfully used ERK-nKTR to examine ERK activity dynamics *in vivo*, and showed that continued Egfr hyperactivation in wing disc clones raises the overall level of ERK activity as compared with control, but does not result in any detectable change to the kinetics.

### Heterogeneity in ERK activity among retinal basal glia reflects their direction of migration

Finally, we asked whether ERK-nKTR could be used in migrating cells with complex morphology to measure EKR activity during development. We focused on the retinal basal glia (RBG), which migrate from the optic stalk into the eye imaginal disc, starting in the third larval instar (Fig. 5A). Upon reaching the disc, RBG are exposed to the FGF ligand Thisbe, expressed by photoreceptors, which induces RBGs to initiate differentiation and wrapping of photoreceptor axons (Franzdóttir et al., 2009). Additionally, they are exposed to the EGF Spitz, also derived from photoreceptors, which induces the glia to express secondary signals involved in patterning the photoreceptor target field, the lamina, in the optic lobe (Fernandes et al., 2017). We labelled control larvae with an antibody against dpERK and observed heterogeneity in ERK activity among RBG in the optic stalk (Fig. 5B), such that some glia displayed high levels of dpERK (Fig. 5B, arrowhead), while others had low levels (Fig. 5B, arrow).

**Figure 5.**
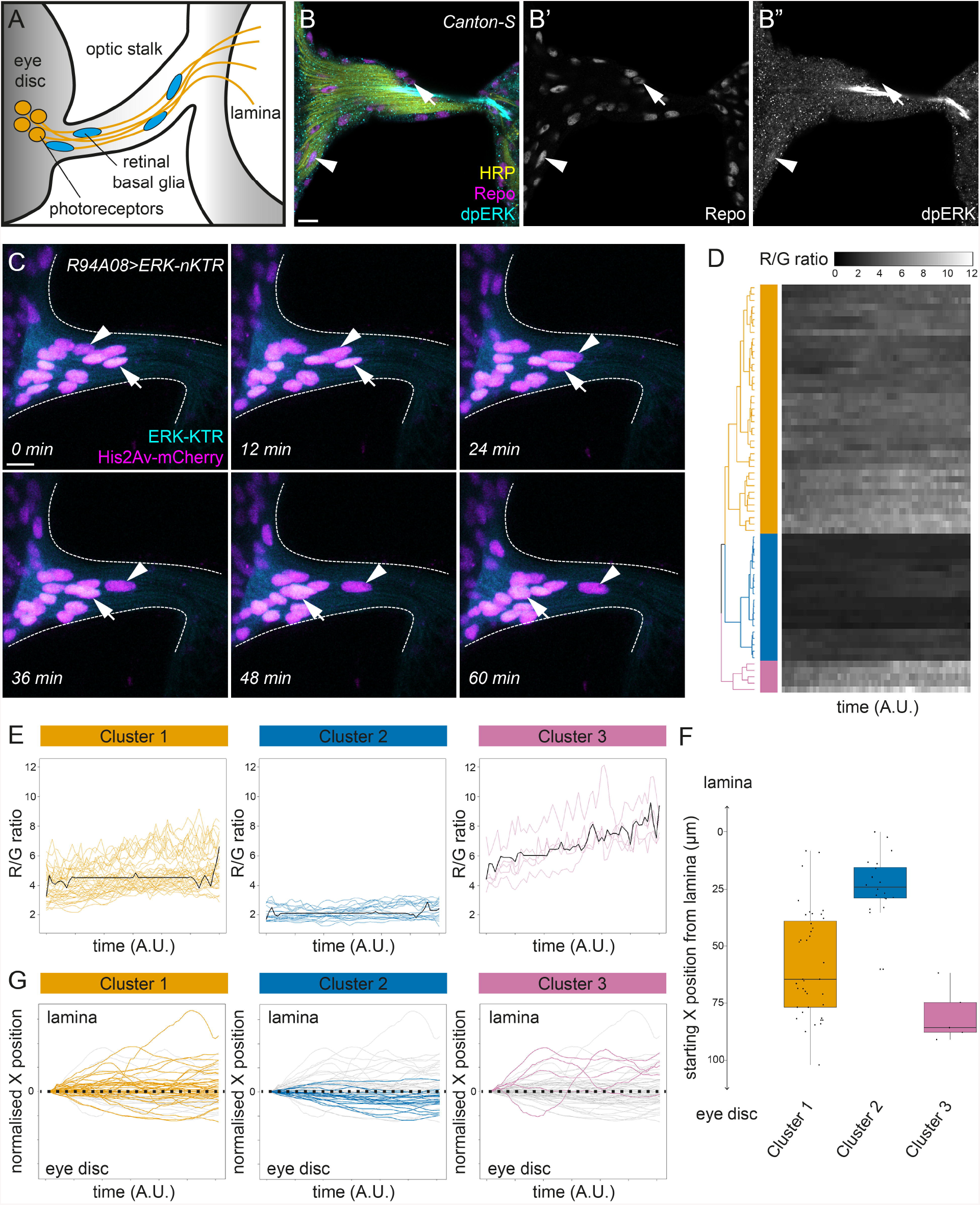
Heterogeneous ERK activity among retinal basal glia reflects their direction of migration. (A) Schematic of the developing Drosophila visual system. Beginning in the third instar larval stage, retinal basal glia (RBG, blue) migrate from the optic stalk into the eye disc. Upon reaching the eye disc, RBG are exposed to cues that induce glial differentiation and wrapping of photoreceptor axons (yellow) that extend down the optic stalk towards the lamina in the optic lobe. (B) dpERK staining (cyan, B”) in control animals, with Repo (magenta, B’) marking RBG and HRP (yellow) labelling photoreceptor axons. Repo-positive RBG are visible in the optic stalk and at the base of the eye imaginal disc, displaying heterogeneous levels of ERK (arrowhead indicates cell with high ERK activity, arrow indicates cell with low ERK activity). (C) Individual timepoints from a time-lapse movie of a developing visual system where ERK-nKTR was expressed in RBG with the *R94A08*-*Gal4* driver. Individual nuclei can be clearly seen migrating towards the lamina (arrowhead) or towards the eye imaginal disc (arrow) over the course of the experiment. (D) Heatmap showing the ERK-nKTR R/G ratios in RBG over time with each row corresponding to a single tracked cell. Traces were hierarchically clustered using dynamic time warping, resulting in three clusters being identified, which were colour-coded in yellow (Cluster 1), blue (Cluster 2), and pink (Cluster 3) in the dendrogram (N = 64 cells across three movies). Time-series were of varying lengths and were thus interpolated for visual representation. (E) Plots showing traces for the ERK-nKTR R/G ratios for cells from each cluster, as colour-coded in panel D, and the cluster average trajectories (black line). (F) Plot showing the starting positions along the X-axis of individual cells from each cluster using the colour code in panel D. Starting positions were normalised to compare samples, such that distances are given relative to the cell closest to the lamina. Boxes outline the interquartile range and the line indicates the median. (G) Plots showing the physical trajectories of cells of each cluster over time, using the colour code in panel D. Trajectories were normalised to the cell’s starting position, as demarcated with a black dashed line, to visualise the direction of travel. Movement above the starting position corresponds to migration towards the lamina, whereas movement below the starting position corresponds to movement towards the eye disc. Scale bar = 10 μm.

To understand the source of this heterogeneity, we conducted time-lapse imaging of ERK levels in wild-type RBG by expressing ERK-nKTR with the RBG-specific driver, *R94A08*-*Gal4* (Movie S3). Over the course of 1.5-3 hours of imaging, we observed RBG nuclei labelled with His2Av-mCherry migrating along the optic stalk in both directions (Fig. 5C, arrowhead indicates cell moving away from eye disc, arrow indicates cell moving towards eye disc). ERK-KTR levels in the nucleus varied significantly between cells, similar to our observations of dpERK immunohistochemistry in fixed samples (compare Fig. 5B with Fig. 5C). We measured ERK-nKTR R/G ratios over time in 64 cells (from 3 independent movies) and clustered the traces using dynamic time warping analysis, as above. This analysis grouped the traces into three clusters, corresponding to cells with high and increasing, intermediate, and low ERK-nKTR R/G ratios (Figs 5D,E and S5C), consistent with our observations in fixed tissues that RBG display varying levels of ERK activity. We asked whether the differences in ERK activity in RBG could be explained by their position or direction of travel; in other words, whether cells with low ERK were those that had not yet been exposed to photoreceptor-derived ligands for the ERK pathway and thus were still ‘naïve’. We plotted the starting point of cells in each cluster, revealing that RBG with low ERK levels were found in the optic stalk closer to the lamina (Fig. 5F, blue), while RBG with higher levels were located closer or within the eye disc (Fig. 5F, pink). Moreover, cells with high ERK-nKTR R/G ratios migrated primarily towards the lamina (Cluster 3 in Fig. 5G), while cells with low ERK-nKTR R/G ratios migrated towards the eye disc (Cluster 2 in Fig. 5G). Thus, analysis of ERK activity in live RBG using ERK-nKTR suggests that the heterogeneity in ERK activity among RBG is due to their developmental state: cells that have not yet reached the eye disc and are undifferentiated have low levels of ERK signalling, whereas glia that have interacted with photoreceptors in the eye disc have high levels of ERK activity and maintain these high levels even when they migrate back down the optic stalk towards the optic lobe.

## Discussion

In order to fully understand how intercellular communication affects cell fate, it is critical to consider the temporal dynamics of signalling. Here we report the adaptation of the KTR technology for capturing ERK activity dynamics in Drosophila. Although initially developed for use in mammalian cell culture (Regot et al., 2014), ERK-KTR has since been successfully adapted in *C. elegans*, zebrafish and mouse and has enabled visualisation of ERK kinetics in a longitudinal manner (de la Cova et al., 2017; Mayr et al., 2018; Okuda et al., 2021; Pokrass et al., 2020; Simon et al., 2020). Combining the powerful advantages of Drosophila genetics with tissue-specific expression using the Gal4/UAS or Q systems means that Drosophila ERK-KTR is a versatile tool that will enable precise dissection of the dynamics of ERK signalling in a variety of physiological contexts. We show that ERK-KTR reports on endogenous ERK activity levels and that it responds to genetic manipulations that increase or decrease ERK activity in larval or adult fly tissues. Additionally, we generated and validated stocks carrying a construct encoding ERK-KTR and a tagged Histone, enabling measurements of ERK-KTR levels using only the nucleus, greatly facilitating live analysis and use in cells where the cytoplasm may not be easy to outline.

In our experiments, under conditions of constitutive Egfr activation for a prolonged period during development, ERK levels are increased without changes in the temporal patterning. This contrasts with observations in cell culture, where ligands can be applied in an acute manner. Previous work has shown that, in rat PC-12 cells, EGF stimulation resulted in a transient increase in ERK activity, whereas NGF stimulation led to a sustained ERK response (Marshall, 1995; Santos et al., 2007). Mimicking the dynamics of NGF-induced ERK activity by supplying repeated EGF pulses resulted in cell differentiation, a phenotype associated with NGF stimulation (Ryu et al., 2015). This implies that the different cell behaviours induced by different ligands are caused by differential temporal profiles of ERK activity. Though our experiments are more representative of the manipulations usually carried out *in vivo*, we have not studied the effect of hyperactivating other RTKs for direct comparison of ERK dynamics. Thus, whether temporal patterns of signalling influence phenotypic outcomes *in vivo*, or whether these outcomes simply depend on the levels of pathway activation remains to be determined. Alternatively, a combination of the two may be at play. More recently, Johnson and Toettcher (2019) used optogenetics to manipulate ERK activity in the Drosophila embryo and found that the cumulative dose of ERK signalling determines whether a cell adopts an endodermal or an ectodermal fate, irrespective of the pattern of activity. While this shows that cells can integrate the amount of signal received over time to induce different fates at different thresholds, whether the same mechanism explains how different ligands produce different outputs through the ERK cascade remains to be determined. The RBG provide an excellent model in which to ask this question, as two different ligands from the photoreceptors (EGF and FGF) activate ERK through different receptors in the glia, leading to different outcomes (Fernandes et al., 2017; Franzdóttir et al., 2009)). Whether these outcomes are dependent simply on total activity thresholds or encoded by different kinetics of activation could be assessed using ERK-KTR to visualise ERK activity in RBG exposed to EGF, FGF, or both, as in wild-type animals.

Here we instead sought to understand the biological meaning of heterogeneity in ERK activity between RBG. As our ability to interrogate individual cells increases, for instance through single-cell sequencing, it is becoming more important to understand what constitutes a meaningful difference in cell type, versus a cell state, or simply noise. Our live imaging of ERK-KTR show that the levels of ERK activity in RBG corresponds to their direction of migration. This suggests that ERK levels can be used to monitor the developmental stage of glia, with naïve RBG, which have not yet been exposed to photoreceptor-derived signals and having low ERK activity, and maturing glia migrating in the opposite direction experiencing high ERK activity. Thus, in the case of RBG, heterogeneity in ERK levels reflects the developmental state of the glia – a conclusion which would not have been apparent by looking at snapshots of ERK activity at specific timepoints.

Beyond ERK signalling, as the modular design of KTR technology easily lends itself to implementation for other kinases (Regot et al., 2014), the tool described here should, in principle, be useful for achieving a fuller understanding of different kinase activity dynamics.

## Materials and Methods

### Fly stocks and husbandry

Fly stocks and crosses were raised on standard cornmeal food at 25□. For experiments in the testis, crosses were raised at room temperature, and freshly eclosed males of the correct genotype were shifted to 29□ for two days prior to dissection. Flip-out clones were induced by heat shocking larvae in a water bath at 37□ for 6 or 10 minutes two days post-egg laying for fixed and live imaging experiments, respectively. Early third instar larvae were selected for dissection four days post-egg laying.

We used the following fly stocks: *Oregon-R, Tubulin* (*Tub*)-*Gal4*; *act-FRT-STOP-FRT-Gal4, UAS-lacZ* (gift of Y. Mao); *traffic jam* (*tj*)-*Gal4* (Kyoto DRGC#104055); *C587*-*Gal4* (gift of R. Lehmann); *R94A08*-*Gal4* (BDSC#40673); *UAS*-*Ras*^*V12*^; *UAS*-*DN-Egfr*; *UAS*-λ*Top*; *UAS*-*Hop* (gift of E. Bach).

### ERK-KTR sequence and stock generation

ERK-KTR sequences derived from the sequence published by Regot *et al*. (2014) (Addgene#59138) were codon-optimised for Drosophila and synthesised by ThermoFisher Scientific GeneArt, and flanked by EcoRI and SalI sites. These were used to subclone the sequence into the EcoRI and XhoI sites of pUASt-attB (gift of W. Wood) and QUAS-attB (gift of N. Tapon). For His2Av-mCherry, the His2Av and linker sequences used by Clarkson and Saint (1999) were synthesised and fused to mCherry. To generate ERK-nKTR, ERK-KTR and His2Av-mCherry were separated by a T2A sequence and a GRAGGS linker, following de la Cova *et al*. (2017). Transgenic flies were generated by injection of the plasmids into flies carrying either attP40, attP2, attP1, or attP64 landing sites and integrated using ΦC31 integrase. Injections were carried out by BestGene Inc.

UAS-ERK-KTR, UAS-ERK-nKTR, and QUAS-ERK-nKTR stocks, together with UAS-His2Av-mCherry, QUAS-His2Av-mCherry, UAS-His2Av-EGFP, and QUAS-His2Av-EGFP stocks have been deposited and are available from the Bloomington Drosophila Stock Center, Bloomington, Indiana, USA.

### Immunohistochemistry

Following dissection, samples were fixed in 4% paraformaldehyde in PBS for 15 minutes, then washed twice for 30 minutes in PBS with 0.5% Triton X-100. Samples were blocked in PBS with 0.2% Triton X-100 and 1% Bovine Serum Albumin (PBTB), before adding primary antibodies and incubating overnight at 4□. Samples were washed twice for 30 minutes in PBTB, then incubated with secondary antibodies for 2 hours at room temperature before washing twice more for 30 minutes in PBS with 0.2% Triton X-100. Samples stained with anti-dpERK were dissected and fixed in a buffer containing 10mM Tris-HCl (pH 6.8); 180 mM KCl; 50 mM NaF; 10 mM NaVO_4_; and 10 mM β-glycerophosphate as previously described (Schulz et al., 2002). Samples were mounted on slides with Vectashield medium (H-1000, Vector labs).

We used the following primary antibodies: chicken anti-GFP (1:500, Aves Labs GFP-1010); rabbit anti-phospho-ERK (1:100, Cell Signaling 9101S); guinea pig anti-Tj (1:3000, gift of D. Godt); AlexaFluor405 conjugated goat anti-HRP (1:100, Jackson ImmunoResearch). Mouse anti-Repo-8D12 (1:20) and mouse anti-Dlg-4F3 (1:20) were obtained from the Developmental Studies Hybridoma Bank and were deposited by C. Goodman. The following nuclear dyes were used: DAPI (1:1000, Sigma-Aldrich D9542); To-Pro-3 (1:500, Invitrogen T3605). Secondary antibodies were obtained from Jackson ImmunoResearch or Invitrogen and used at 1:200.

### Imaging and image processing

Samples were imaged using a Leica Sp8, a Zeiss LSM800, or Zeiss LSM880 confocal microscope. Adult wing images were acquired on a Nikon SMZ1270 microscope fitted with a Moticam X camera. To measure the ERK-KTR C/N ratios and ERK-nKTR R/G ratios, nuclei were manually segmented on ImageJ, using the freehand selection or selection brush tools, at the plane with the largest nuclear diameter as delineated by either DAPI, To-Pro-3, Tj, or His2Av-mCherry fluorescence. For fixed eye imaginal discs, the cytoplasmic mClover intensity was measured by expanding the nuclear selection by 3 pixels. For wing imaginal discs and ovarioles, cytoplasmic mClover intensity was determined using Dlg staining to outline cells. To ensure that dpERK intensities could be compared across samples, min-max normalisation of nuclear dpERK intensity was performed for each disc and ovariole.

Live imaging experiments were performed with samples embedded in agarose gel as described in Bostock *et al*. (2020). For wing discs, confocal stacks were acquired at two-minute intervals with a z-step of 1 μm. For RBG movies, confocal stacks were acquired every three to four minutes with a z-step size of 2 μm. Movies were processed with the Bleach Correct plugin on ImageJ. Quantification of nuclear mCherry and mClover intensity was carried out on ImageJ by manually segmenting nuclei at the plane with the largest nuclear diameter, identified using the His2Av-mCherry channel, across timepoints and saved as regions-of-interest (ROI) on ROI Manager. For RBG tracking, XY coordinates of cells were obtained by measuring the centre of mass of each ROI. Representative movies were prepared using Adobe Premiere Pro.

### Data analysis

Statistical tests were performed using GraphPad Prism. Significance for clonal perturbation experiments was computed using one-way ANOVA followed by Holm-Šidák’s multiple comparisons test to determine differences between multiple samples (Figs 2I,J and S3). Two-tailed, unpaired Student’s t-test was performed for adult wing size (Fig. S2C) and for mean ERK-nKTR R/G ratios in wing disc live imaging experiments (Fig. 4C). ERK-KTR and ERK-nKTR localisation as a function of normalisation dpERK levels were fitted to asymmetric sigmoidal curves (Figs 1F,G and 3G,H); the R-squared value for each is indicated on the graph. ERK-nKTR^AA^ and ERK-nKTR^EE^ R/G ratios as a function of normalised dpERK levels were fitted to simple linear regressions (Fig. 3G), as were ERK-KTR C/N ratios as a function of total ERK-KTR levels (Fig. S1). Plots show mean and 95% confidence intervals. Dotted lines in violin plot for wing disc live imaging experiments (Fig. 4C) shows median, upper and lower quartiles. Box plot for RBG live imaging experiments (Fig. 5F) shows median and interquartile range of starting positions for each cluster.

For clustering ERK activity trajectories, the *dtw* package on R was used to obtain distance matrices that were then processed by hierarchical clustering. The optimal number of clusters was determined by an elbow plot (Fig. S5). Heatmaps, percentage bar charts, and trajectory plots were generated on R Studio. Time series from live imaging of RBG were of variable lengths and therefore interpolated to span the entire time axis for visual representation. Time series averages were generated using DTW barycenter averaging using the Incremental Calculation of Dynamic Time Warping (*IncDTW*) package and interpolated on R. For plotting RBG trajectories, as the samples were oriented such that the eye discs and lamina were located at opposite ends of the image (left and right, respectively), position along the X-axis was used to represent the direction of migration.

## Acknowledgements

We are grateful to members of the Amoyel and Fernandes labs for feedback and discussions. Many thanks in particular to Pablo Araguas Rodriguez for insightful discussions on coding and data analysis. We thank members of the fly community for stocks and reagents.

